# Wavefront-sensorless adaptive optics with a laser-free spinning disk confocal microscope

**DOI:** 10.1101/2020.07.28.225607

**Authors:** Syed Asad Hussain, Toshiki Kubo, Nicholas Hall, Dalia Gala, Karen Hampson, Richard Parton, Mick A. Phillips, Matthew Wincott, Katsumasa Fujita, Ilan Davis, Ian Dobbie, Martin J. Booth

**Affiliations:** Department of Engineering Science, University of Oxford, Parks Road, Oxford OX1 3PJ, United Kingdom; Department of Applied Physics, Osaka University, Osaka 565-0871, Japan; Micron Advanced Bioimaging Unit, Department of Biochemistry, University of Oxford, South Parks Road, Oxford OX1 3QU, United Kingdom; Department of Biochemistry, University of Oxford, Oxford, United Kingdom

## Abstract

Adaptive optics is being applied widely to a range of microscopies in order to improve imaging quality in the presence of specimen-induced aberrations. We present here the first implementation of wavefront-sensorless adaptive optics for a laser-free, aperture correlation, spinning disk microscope. This widefield method provides confocal-like optical sectioning through use of a patterned disk in the illumination and detection paths. Like other high-resolution microscopes, its operation is compromised by aberrations due to refractive index mismatch and variations within the specimen. Correction of such aberrations shows improved signal level, contrast and resolution.

## Introduction

Three-dimensionally resolved fluorescence imaging is a vitally important method in biomedical microscopy. Several methods of optically sectioning widefield microscopes are available as complementary approaches to the widely used laser point-scanning confocal microscopes. Many of these methods are, in essence, parallelised versions of the confocal laser scanning microscope, based upon the spinning Nipkow-Petran disk [1], [2]. Another approach uses lamp or light emitting diode (LED) based illumination in conjunction with a patterned disk placed in the illumination and imaging paths [3]–[5]; this optical sectioning approach relies upon correlations between the patterns used in the illumination and detection paths. All of these methods suffer from problems caused by system and specimen-induced aberrations, which arise from the inhomogeneous refractive index structure of thick specimens. As in all microscopes, such aberrations affect imaging quality, through loss of resolution and contrast.

Adaptive optics (AO) has been introduced into microscopes to correct aberrations and restore image quality [6]–[9]. An adaptive element, such as a deformable mirror or liquid crystal spatial light modulator, is built into the microscope in order to compensate the aberrations introduced by the specimen [6]. Different AO correction methods have been implemented for different microscope modalities. A common approach is to use the so-called wavefront-sensorless AO methods (or “sensorless AO” for short) in microscopes [9]. These methods estimate the best correction for the microscope by inferring the input aberrations from a sequence of images taken when a sequence of predetermined aberrations is applied with the adaptive element. The optimal design of such AO methods depends strongly on the nature of aberrations and the image formation process of the microscope.

This form of sensorless AO has been implemented previously in a Nipkow-Petran type widefield microscope [10]–[12], which has imaging properties closely related to the confocal microscope. It has also been demonstrated in widefield and super-resolution structured illumination microscopes [9]. However, such aberration correction has not yet been shown in the correlation disk type microscopes, which have considerably different image formation processes. In this paper, we demonstrate a practical implementation of AO in this microscope, covering both hardware and control aspects, by incorporating a custom deformable mirror based AO unit with a Clarity microscope module (Aurox Ltd, United Kingdom).

### Imaging process and effects of aberrations

The correlation disk microscope is fundamentally a widefield fluorescence incoherent imaging system. A disk is placed in an image plane in the common illumination and imaging path, Fig. 1. The disk is imprinted with a binary pattern that either reflects or transmits the incident light. In the illumination path, the light generated by the LED source impinges upon the disk; part of this light is reflected by the pattern and discarded; the other part passes through the pattern and illuminates the specimen. The patterned illumination selectively excites fluorescence in the focal plane of the specimen. It is important to note that this pattern only appears near the focal plane and in out-of-focus planes the illumination is uniform. The fluorescence emitted by the specimen is imaged back onto the disk, whereby that part of the light in positions corresponding to the brightly illuminated parts of the focal plane passed through the pattern on the disk. The remaining emission is reflected by the disk into a separate beam path. The paths are arranged such that both images (transmitted through and reflected off the disk pattern, respectively) fall onto two halves of the camera chip. The two images are aligned with the help of calibration pattern inside the box and shown in yellow in Fig. 1. The disk is rotated at a speed much faster than the camera frame rate in order to average out the appearance of the pattern on the camera images.

**Fig. 1.**
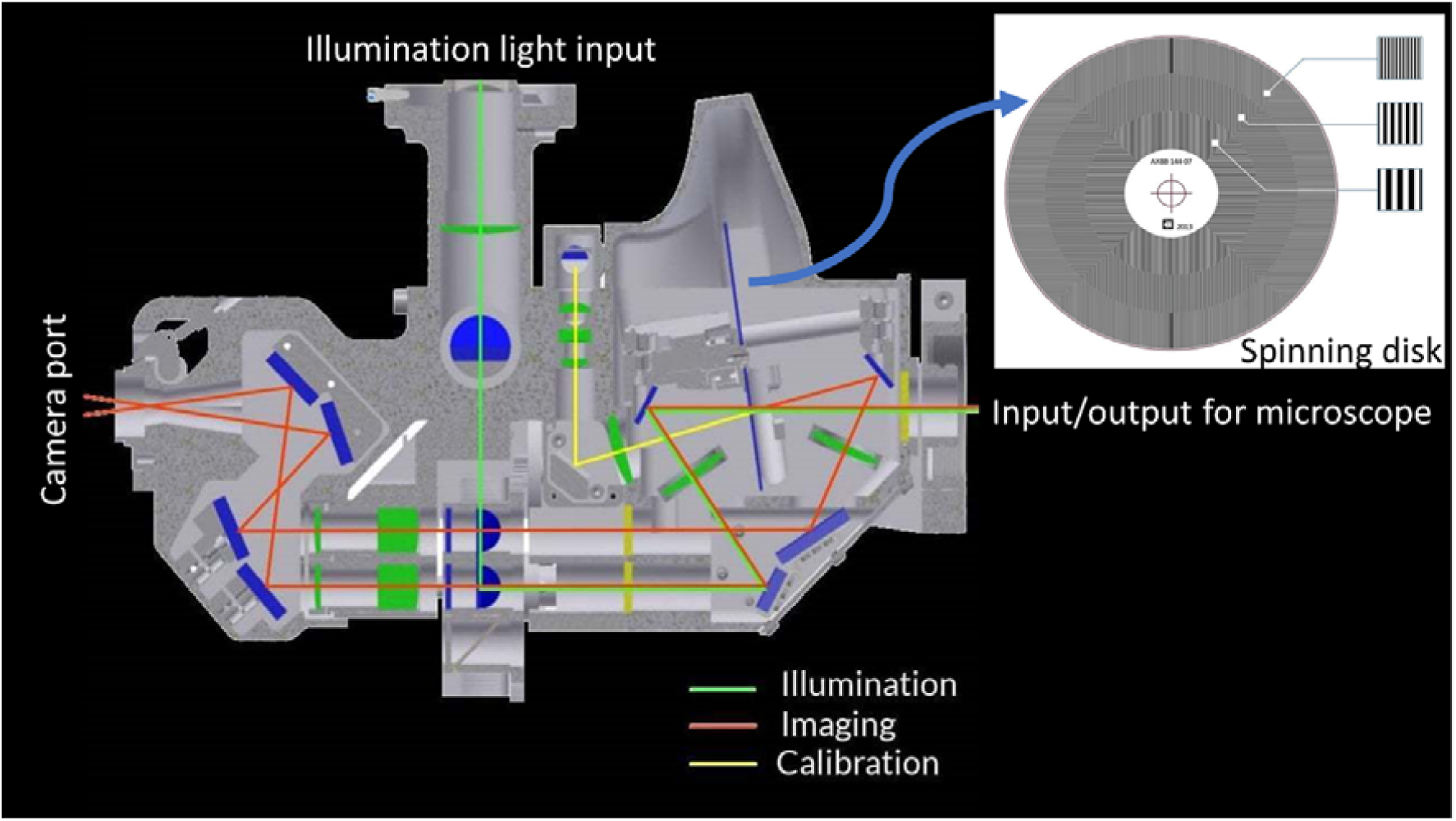
Cross sectional view of the Aurox clarity module, which is placed between the microscope port and the camera and light source. The spinning disk inside the system includes three patterns with different spacings for different sectioning strengths. (Modified with permission from Aurox Ltd., Copyright Aurox Ltd.).

It has been shown [3], [4] that the disk-transmitted image *I*_*T*_ is in effect a sum of a conventional microscope image *I*_*conv*_ and a sectioned image *I*_*sect*_, whereas the reflected image *I*_*R*_ is the conventional image minus the sectioned image:

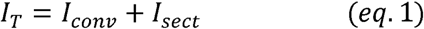

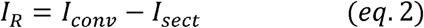

Following appropriate image registration processes, it is thus possible to extract the conventional and sectioned images simultaneously as

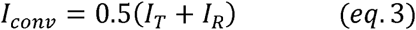

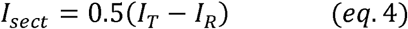

The exact nature of the sectioned image depends upon the properties of the microscope optics, including objective lens numerical aperture (NA), and of the disk pattern. The Aurox Clarity disk patterns consist of parallel, equally spaced stripes with unit mark-space ratio. Different patterns are available on the disk with different stripe spacings, which provide different sectioning ratios, with finer patterns providing stronger sectioning, albeit at lower contrast. This parallels closely the phenomena seen when varying the pinhole size in a point scanning confocal microscope, where a smaller pinhole provides better sectioning, albeit with lower signal level.

When aberrations are present in this microscope, they will affect both the illumination and the imaging paths. On illumination side, the aberrations will cause a blurring of the disk image in the specimen, thus resulting in lower contrast in the structured illumination pattern. On the detection side, the image of the fluorescence distribution onto the disk will be blurred, such that the distinction between the disk-transmitted image and the disk-reflected image becomes less clear. *I*_*T*_ and *I*_*R*_ become weaker; and the signal level (and hence signal to noise ratio) of *I*_*sect*_ is reduced, coupled with compromised resolution – particularly in the axial direction. At the same time, the mean signal level in *I*_*conv*_ is unaffected, although resolution and contrast are reduced. These phenomena mirror those seen in confocal and conventional microscopes, respectively, in the presence of aberrations.

### Experimental system

In normal operation, the Clarity disk module would be attached directly to the microscope side port. In our case, we separated the module from the microscope base (iX71, Olympus, Japan) in order to slot in the AO system. To do this, we created an intermediate image plane into which we could insert the DM (Mirao 52e, Imagine Optics, France), Fig. 2. The additional optics were designed so that the DM was imaged on the objective pupil plane within the microscope. Additionally, the image plane near the camera port of the microscope was reimaged onto the input port of the Clarity module using a 4f lens system. The 4f system used two 300 mm focal length achromatic doublets (Thorlabs, United States) to form a unity magnification telescope. The DM was placed in the intermediate pupil plane of this 4f system. Imaging was performed using an oil immersion microscope objective (PlanApo, Olympus, Japan; 60× magnification, NA 1.4). The focussing depth could be controlled using a piezo z-stage (P-736.ZR1S PI-nano Z Microscope Scanner, Physik Instrumente, Germany) attached to the setup. The stage was able to move with a step size of 0.2nm. Images from both the transmitted and reflected light paths were captured by the camera (Prime BSI, Photometrics, UK), which has pixel size of 6.5μm x 6.5μm; the area of the whole sensor area was 176.89mm^2^ (13.3mm × 13.3mm or 18.8mm diagonal). As described earlier, the two images were directed to the sensor in such a way that the two images were detected separately on the two halves of the sensor. The sectioned (confocal) image was obtained using equation 4. For the study presented here, we used a commercial LED source (PE 300 ultra, CoolLED, United Kingdom) to illuminate the sample. The desired excitation and emission wavelengths were selected by using set of dichroic filter cubes. In our case we used the following four dichroic filters cubes:

- Dichroic filter 1: Excitation 466nm/FWHM 40nm, Emission 525nm /FWHM 45nm
- Dichroic filter 2: Excitation 554nm /FWHM 23nm, Emission 609nm /FWHM 54nm
- Dichroic filter 3: Excitation 578nm /FWHM 21nm, Emission 641nm /FWHM 75nm
- Dichroic filter 4: Excitation 392nm /FWHM 23nm, Emission 447nm /FWHM 60nm

**Fig. 2.**
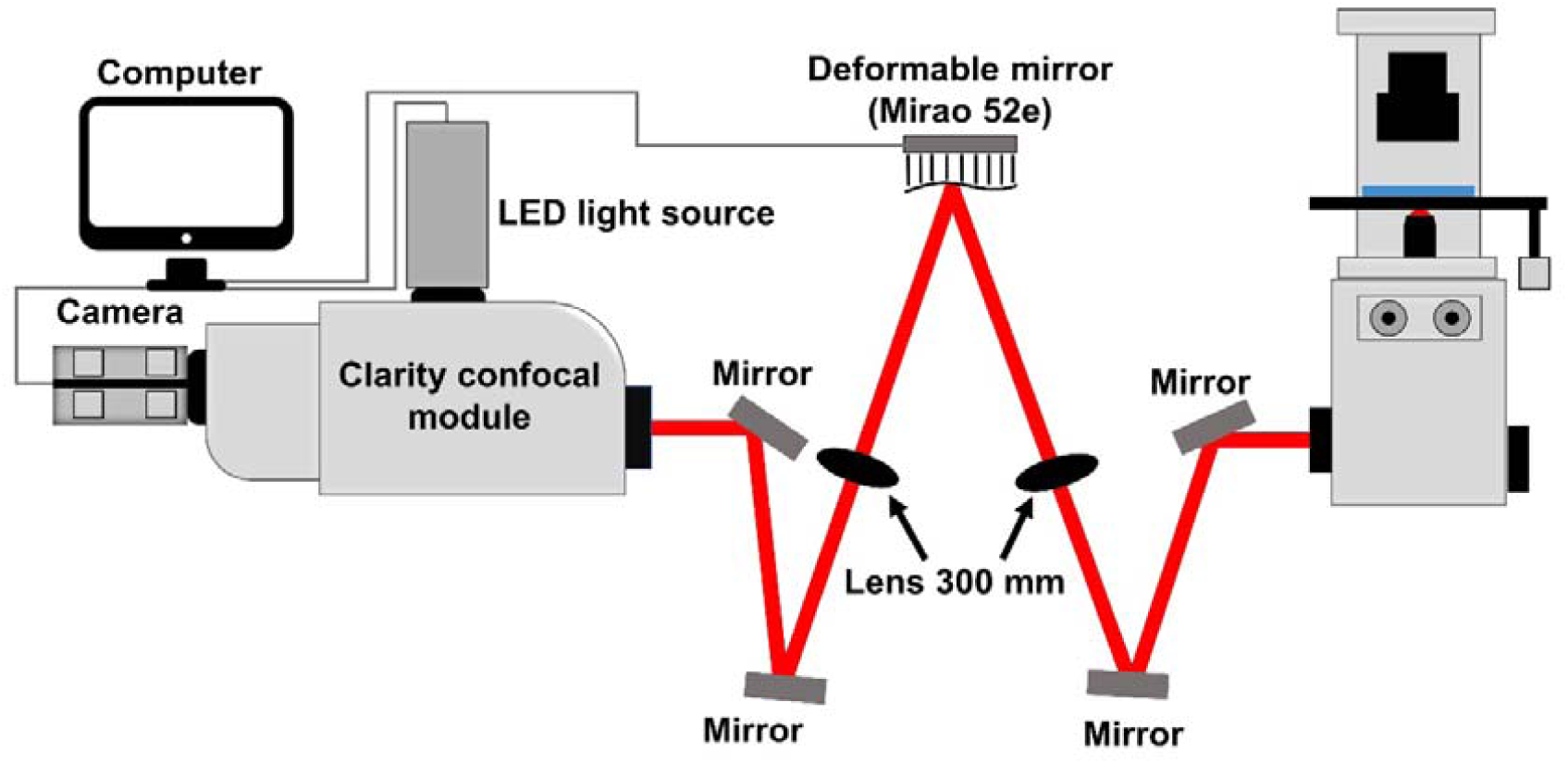
The experimental set-up, consisting of a microscope base, an intermediate imaging system incorporating the deformable mirror, the confocal microscope module, a camera, a light source and a personal computer.

The deformable mirror was calibrated by using a technique based on deflectometry [13], [14]. The control matrix derived at the end of the protocol was used for control of the DM. The whole experimental setup was controlled using a personal computer.

### Adaptive optics strategy

Aberrations were corrected using the deformable mirror via a sensorless AO method, as has already been applied to other adaptive microscopes [9], [15]–[18]. For each aberration mode being corrected, a small number of images are taken with a different amount of the chosen mode applied with the DM. For each image, an image quality metric was determined. These image quality metric values were used to fit a function – in the simplest case, a Gaussian – and the peak of the function is taken to be the optimal correction for the current mode. Different modes were then corrected in sequence.

This method was implemented in Microscope-AOtools, an extension to the Python Microscope hardware control software [19]. Microscope-AOtools offers a suite of image quality metrics and optimisation algorithms. In this case, the image quality metric, *S*, was a weighted sum of the power of all spatial frequencies obtained from the image spectrum. *S* was defined as:

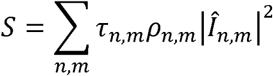

Where *Î*_*n,m*_ is the discrete Fourier transform of the image,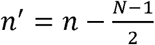 and 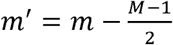 where *n* and *m* are the pixel indices ranging from 0 to *N*-1 and *M*-1 respectively. *N* and *M* are the number of pixels along each image dimension. *τ*_*n,m*_ is the thresholding mask and is defined as:

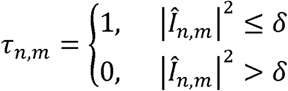

where *δ* is the noise floor threshold defined as:

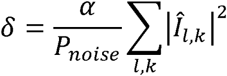

where |*Î*_*l,k*_ |^2^ is the power in the *l* -th, *k* -th spatial frequency where *l* & *k* are the coordinates of the pixels in the regions shown in yellow in Fig. 3a. *P*_*noise*_ is the total number of pixels in the yellow region and *α* is a user defined, arbitrary scaling factor of 1.125. The scaling factor was chosen to be certain we were only using spatial frequency content that was above the noise level.

**Fig. 3a.**
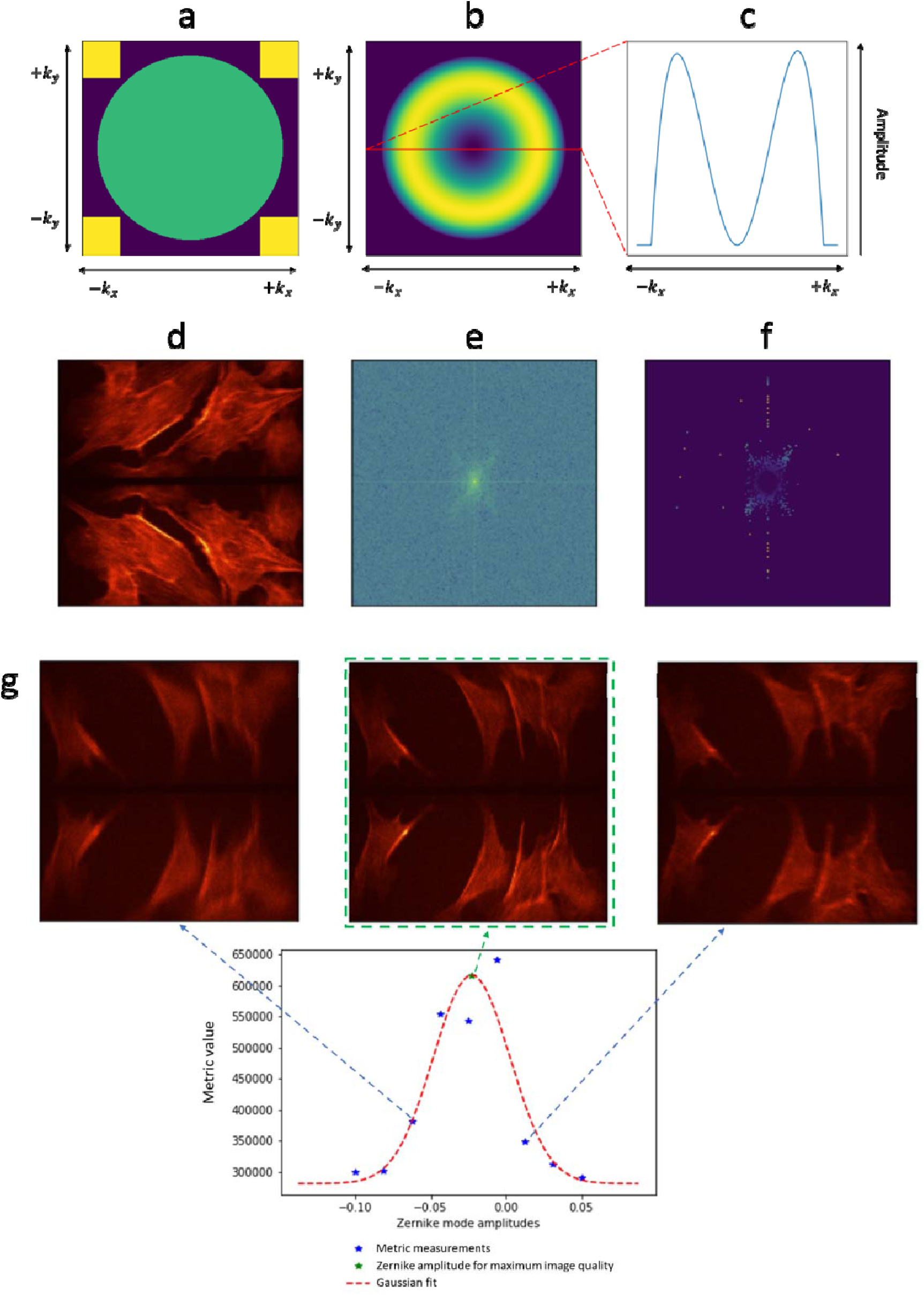
and represent the spatial frequencies corresponding to the and axis respectively. Visualisation of the binary mask,, shown in green and the noise sampling regions for determining, shown in yellow. Fig. 3b shows visualisation of the weighting mask,. Fig. 3c represents cross section of *ρ*_*n,m*_ showing that for calculation of the optimisation metric, the middle-high spatial frequencies are amplified while the very high spatial frequencies (close to 1/θ) are suppressed. Fig. 3d gives an example of a raw image obtained by using bovine pulmonary artery endothelial (BPAE) cells. The image in 3d shows the raw image obtained by the camera; the upper half contains *I*_*T*_ and the lower half *I*_*R*_. Fig. 3e shows the power spectrum of Fig. 3d, |*Î* _*n,m*_|^2^ where *Î* _*n,m*_ is the discrete Fourier transform of Fig. 3d. Fig. 3f shows the power spectrum with both *τ*_*n,m*_ and *ρ*_*n,m*_ applied. This amplifies middle-high spatial frequencies presented in Fig. 3e and thresholds out all spatial frequencies outside of the OTF radius. The image quality metric, *S*, is the sum of all pixels in Fig. 3f. Fig. 3g provides an example showing the variation of image quality metric versus the applied correction of horizontal coma (Zernike mode 8, OSA indexing). A Gaussian function was fitted to the data to estimate the position of the peak, corresponding to the optimal correction. The appropriate aberration correction was then applied to obtain the corrected image.

*ρ*_*n,m*_ is a weighting mask (Fig. 3b) and is defined as:

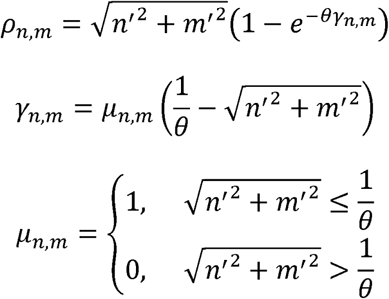

where. *μ*_*n,m*_ is the binary mask shown in green in Fig. 3a and θ is the resolution limit of the microscope given by 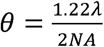. The effect of *ρ* is to amplify the power in the middle-high spatial frequencies of the optical transfer function (OTF), shown in the Fig. 3b. Fig. 3c shows this by taking a cross section in the centre. This makes the metric sensitive to image details that are most affected by aberrations (i.e. structures with mid-high spatial frequency content). The rationale for this choice of optimisation metric is based upon the image formation process of the microscope, which relies upon the high-fidelity reproduction of the structured illumination pattern in the specimen and its imaging back onto the disk. The pattern frequencies are set to be in the mid-range of spatial frequencies covered by the OTF support. The power in low frequencies is barely affected by aberrations and the highest frequencies tend to be dominated by noise. Hence, the enhancement of the mid-range frequencies in the optimisation should provide appropriate feedback for aberration correction.

The metric was calculated using the entire image captured by the camera. Optimisation was performed using the raw image data (rather than the processed sectioned or conventional images), as all of the necessary information for optimisation is contained within the image spectrum of the raw data. Fig. 3d shows an example of a raw image for which we measure the image quality metric. Fig. 3e show the power spectrum, |*Î* _*n,m*_|^2^, of the image and Fig. 3f shows the power spectrum with *τ*_*n,m*_ and *ρ*_*n,m*_ applied.

As described above, to obtain the optimum correction for a specific mode, a number of images (typically 10) were obtained with different amplitudes (typically in the range [−0.06,0.06]) of the Zernike mode applied. The image quality metric was calculated for each image. These values were then fitted to a Gaussian function, the peak of the function obtained and applied to the DM. This process was repeated for each Zernike mode. Fig. 3g shows one such example along with the correction applied and obtained pictures. Finally, all these corrections were performed on one channel by using dichroic filter 1 (excitation 466nm). Aberrations arising from the refractive index structures in the specimen should be corrected for all wavelengths by the deformable mirror, so correction performed on one channel should be valid for all.

## Results

### Aberration correction for different sectioning patterns

The microscope provides three disk patterns of differing spacing that can provide different optical sectioning. We demonstrate here the capabilities of the AO scheme in correcting each of these imaging modes. For all results in this paper, we implemented correction of the low order Zernike aberration modes OSA index 3, 5, 7, 8, and 12, which are oblique astigmatism, vertical astigmatism, vertical coma, horizontal coma and primary spherical, respectively. Fig. 4 shows examples of such aberration correction using a *Drosophila* larva neuromuscular junction (NMJ) sample (see methods for details). In figure 4, we see the synaptic boutons visualised with the Alexa488 fluorophore. In this case, we detected only one channel using dichroic filter 1 (excitation 466nm).

**Fig. 4.**
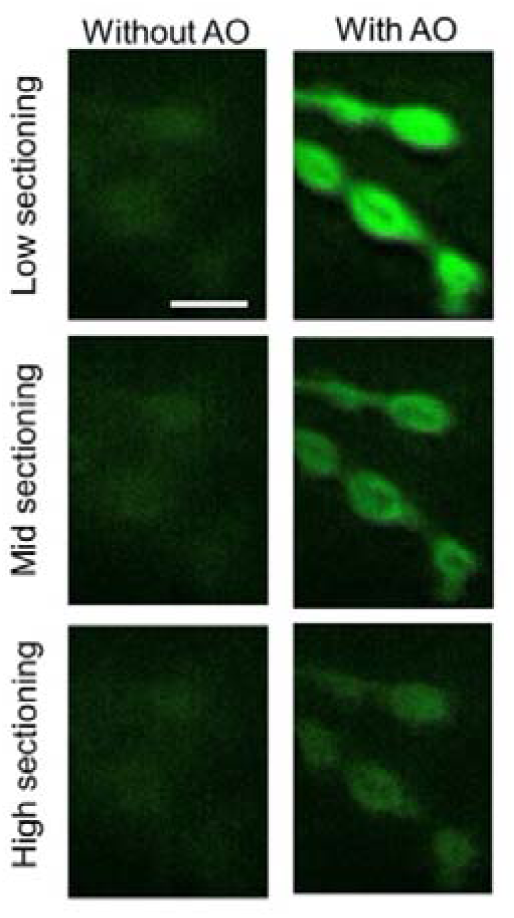
Experimental results when using different settings of the disk. The images were obtained from the NMJ sample. In this case we used only dichroic filter 1 (excitation 466nm). Scale bar 10μm.

The panels on the left labelled “Without AO” show the retrieved section image before aberration correction. Low sectioning using the coarsest disk pattern leads to a thicker optical section; the finest spacing corresponds to a thinner optical section. In all of the “Without AO” cases, the contrast and resolution are poor. This is due to the blurring out of the structured illumination and, hence, a reduction in the difference between the *I*_*T*_ and the *I*_*R*_ images.

On the other hand, after applying the AO correction we obtained the “With AO” images that showed greater contrast and detail. The variation in brightness between the low, medium and high sectioning modes was due to there being less fluorescent marker within the narrower optical sections. The images were taken at the depth of 10μm inside the sample. In each case, this clearly shows that the sensorless AO algorithm functions in all three modes. In the rest of this paper, we present only the results obtained by using the high sectioning mode of the microscope, which is expected to be more sensitive to aberrations.

### Aberration correction varying with depth

Fig. 5 shows images of a *Drosophila* neuro-muscular junction (NMJ; see methods for details) before and after correction at varying focussing depth. These images were taken at depths of 10, 13.5, 26μm inside the NMJ sample. In the figure, we see the synaptic boutons visualised with the Alexa488 fluorophore (see methods). One channel was acquired using dichroic filter 1 (excitation 466nm).

**Fig. 5.**
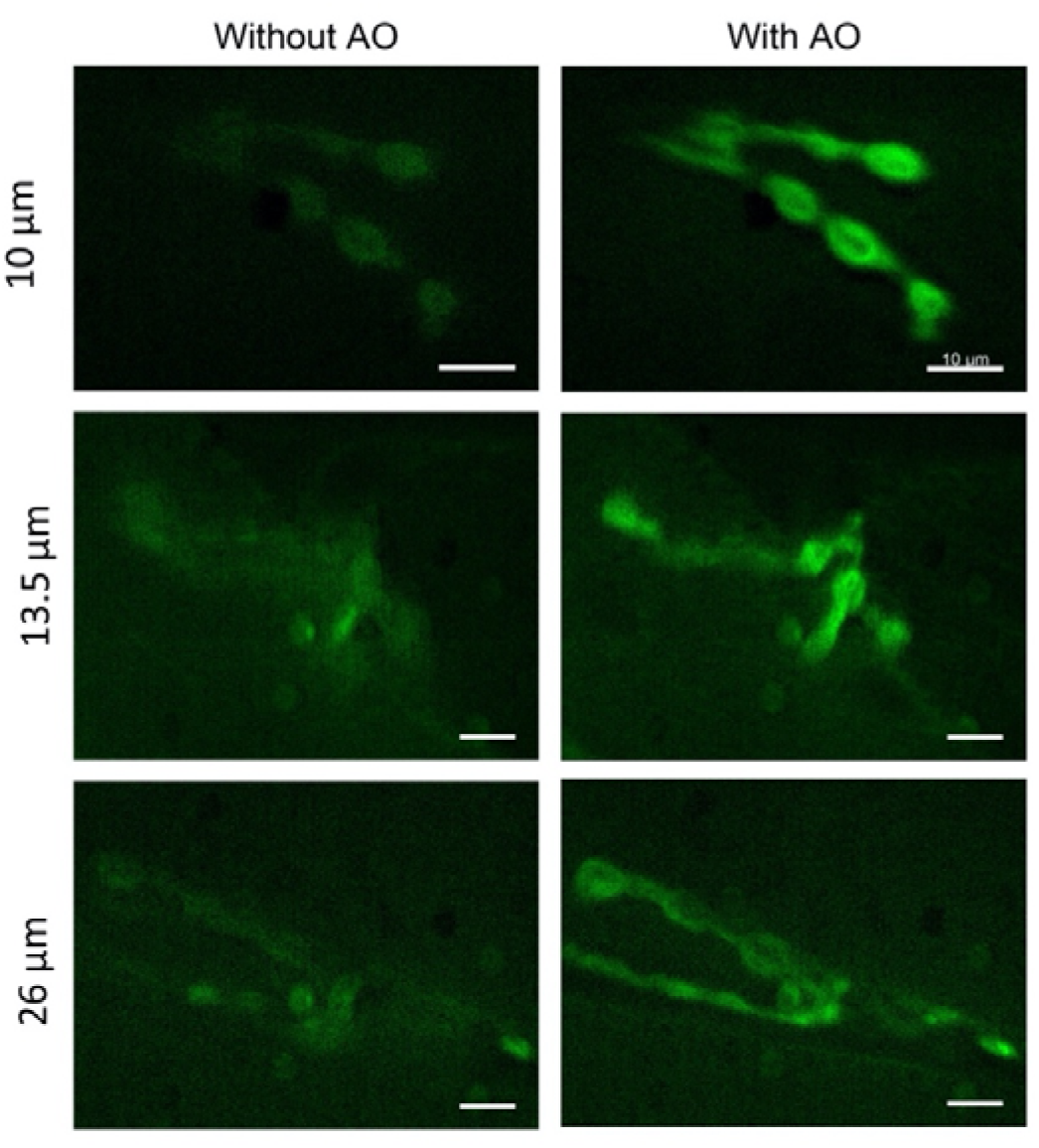
Images taken at depths of 10, 13.5 and 26μm in the NMJ sample when dichroic filter 1 (excitation 466nm) was used. Aberration correction improved imaging quality at all depths. Scale bars 10μm.

At each depth in the sample, the AO correction provided improved contrast and resolution. When focussing to deeper planes into the sample, the signal level decreased even in the corrected images, which may be due to varying marker density, residual aberration or loss of signal through absorption or scattering as the light passes through more tissue.

Fig. 6 shows results obtained for the maximum intensity projections of 20μm z-stacks through an NMJ sample. To gain more insight, we have presented zoomed images of sample regions. We can see in the middle panel that the fine details are only visible when we applied correction. In this case, we were able to see the structure of the fly nerve and synapse better. In the bottom right panel, we can see individual round postsynaptic densities – boutons – which are blurred and not well defined in the bottom right panel.

**Fig. 6.**
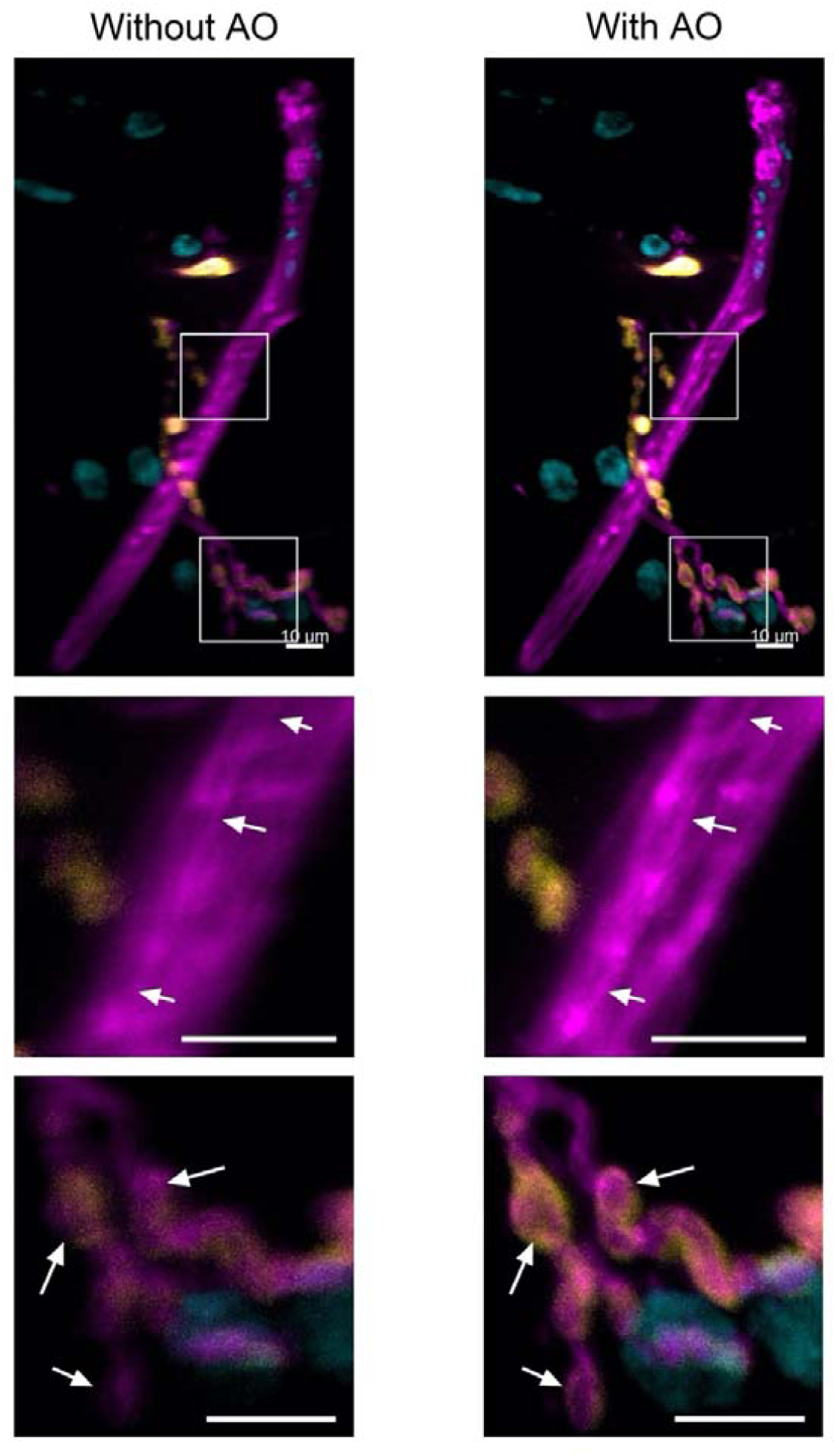
First row are maximum intensity projections of 20μm z-stacks through an NMJ sample. The middle row shows one region of interest in which we can see clear features indicated by arrows that were not visible without AO. The lower row shows a comparison of another region of interest in which individual round postsynaptic densities are clearly visible in the corrected images. Images were taken in three channels using dichroic filters 1, 2 and 4 (excitation 466, 554 and 392nm respectively); the NMJ synapse with the neuron is shown in magenta, postsynaptic density in yellow and the nuclei in cyan Scale bars 10μm.

Fig. 7 shows images of early stage *Drosophila* egg chambers (2-5), approximate sample thickness 100μm. Ovaries were stained for actin in multiple colours using Phalloidin Alexa dye conjugates. Green emission was imaged using dichroic filter 1 (excitation 466nm). Ovarioles were separated out to better visualise individual egg-chambers. Arrows indicate actin labelling associated with ring-canals, pores which connect adjacent nurse cells and the oocyte to adjacent nurse cells the egg chamber. As we can see in the zoomed version, the fine details that were not visible in the non-corrected version become visible in the corrected version. In this case we would like to highlight the information in the lower panel in which the structure appears ring-like after correction but looks filled in for the non-corrected image.

**Fig. 7.**
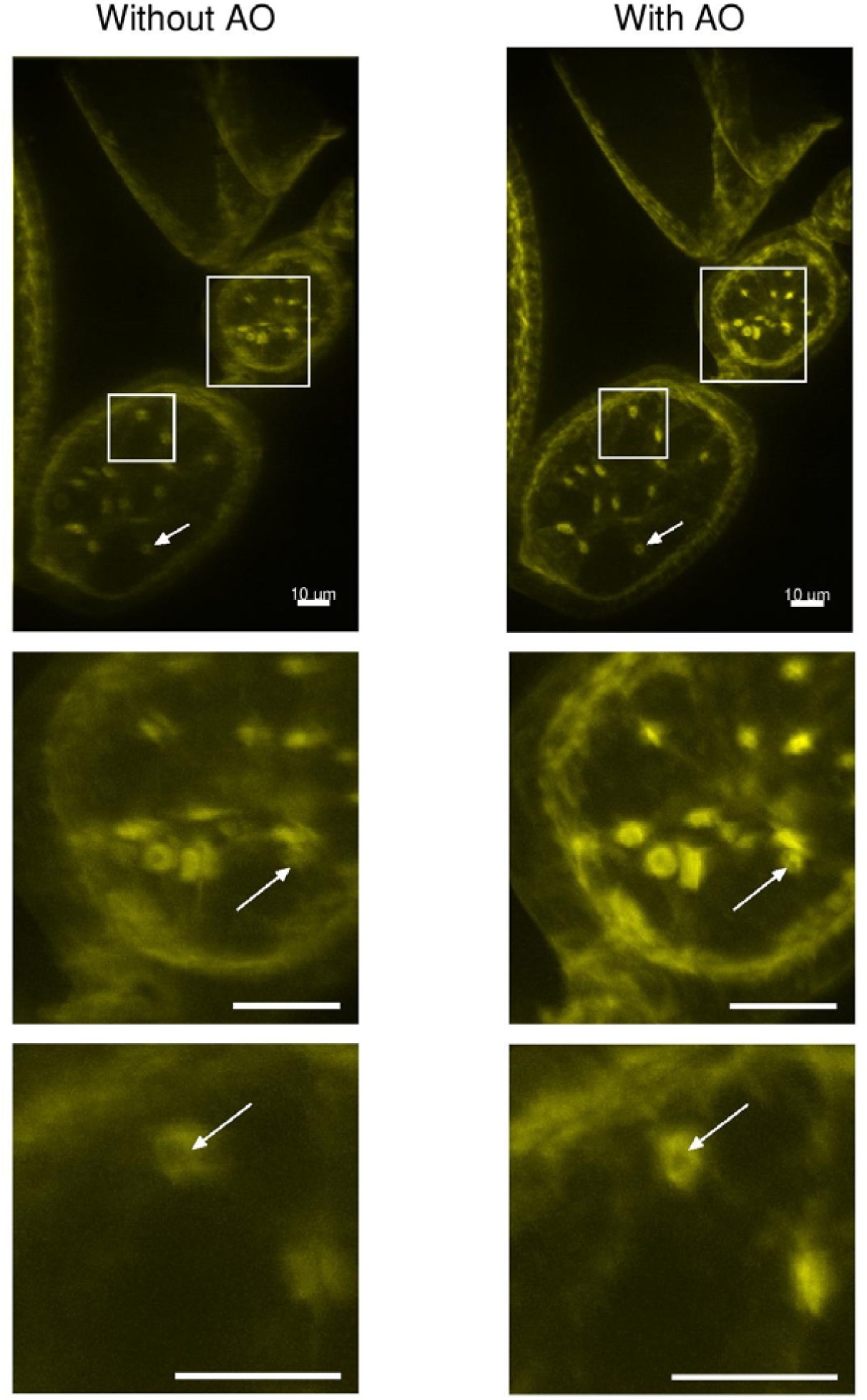
Egg chambers sample imaged without and with AO, first row (dichroic filter 1, excitation 466nm). The middle and bottom represent zoom regions and we can see that small features are clearly visible when AO was applied. Scale bars 10μm.

## Methods

Results were obtained from three samples: bovine pulmonary artery endothelial (BPAE) cells, *Drosophila* NMJ and *Drosophila* egg chambers. The detailed description of the samples is as follows:

### BPAE cells

Cells (F36924, Invitrogen) were fixed and stained with variety of fluorescent dyes. Mitochondria were labelled with red-fluorescent MitoTracker^®^ Red CMXRos (excitation 579nm). Alexa Fluor^®^ 488 was used to stain F-actin (excitation 505nm). Finally, the nuclei were stained with blue-fluorescent DAPI (excitation 358nm).

### NMJ

The samples were prepared by following the protocol presented in Ref.[20]. 3^rd^ instar *Drosophila melanogaster* larvae (Oregon-R strain) were dissected in HL3 buffer with 0.3mM Ca^2+^ to prepare a so-called larval fillet. After this, larvae were fixed with paraformaldehyde and blocked using BSA. Larvae were stained overnight with HRP (conjugated to Alexa568 fluorophore, visible when dichroic filter 2 (excitation 554nm) was used) to visualise the neurons, and primary mouse antibody against DLG - discs large – to visualise the postsynaptic density. The next day, the larvae were counterstained with secondary antibody to detect the DLG (donkey anti-mouse conjugated to Alexa488 fluorophore, visible when dichroic filter 1 (excitation 466nm) was used), as well as DAPI to visualise the nuclei (visible when dichroic filter 4 (excitation 392nm) was used). The larvae were then washed and mounted in 65% vectashield (this dilution of vectashield is compatible with the oil immersion lens that was used).

### Egg chambers

Ovaries were isolated from *Drosophila melanogaster* adult females straight into fixation medium and fixed in 4% PFA (prepared fresh from a stock solution of 16% paraformaldehyde, methanol free ultra - pure EM grade, Polysciences) in Cytoskeleton Buffer [21], [22] for 10 minutes without added detergent [23]. Ovaries were then heptane permeabilised and post fixed in 4% PFA in PBS for not more than 15 minutes, then washed three times with PBS / TX-100, 0.05%. Actin was labelled with Phalloidin Alexa 488 (Phalloidin: stock 200 units/ml in Methanol, 6.6 μM) at between 5 and 20 μl stock in 200 μl for at least 45 min at RT (or O/N at 4°C). This was visible when dichroic filter 1 (excitation 466nm) was used. Labelled ovarioles were washed three times with PBS TX-100, 0.05%, and then finally with PBS before being pre-incubated at RT for at least 3 h in 70% Vectashield. This was removed and replaced with fresh 70% Vectashield prior to samples being mounted under a coverslip raised on two thin strips of double-sided sticky tape to avoid flattening[24]. The edges of the coverslips were double varnished, and slides were stored at 4°C until use.

## Conclusion

We have introduced a sensorless AO correction scheme that is applicable to correlation disk microscopy. Aberration correction has been achieved by the optimisation of an image quality metric that is designed to be sensitive to spatial frequencies in the mid-range of the OTF pass band, as these are the most important for effective imaging in this system. The scheme was shown to be suitable for different sectioning modes and fluorescence channels, as in all cases it was able to increase signal levels and reveal image details that were otherwise blurred. The AO methods should be widely applicable for other imaging applications, extending the capabilities of these microscopes for deep tissue imaging.

